# Foveal vision at the time of microsaccades

**DOI:** 10.1101/2021.01.26.427903

**Authors:** Naghmeh Mostofi, Janis Intoy, Michele Rucci

**Affiliations:** Department of Psychological and Brain Sciences Boston University.; Graduate Program for Neuroscience, Boston University.; Department of Brain & Cognitive Sciences University of Rochester, 310 Meliora Hall, Rochester, NY 14627.; Center for Visual Science, University of Rochester, 310 Meliora Hall, Rochester, NY 14627.

## Abstract

Humans use rapid eye movements (saccades) to inspect stimuli with the foveola, the region of the retina where receptors are most densely packed. It is well established that visual sensitivity is generally attenuated during these movements, a phenomenon known as saccadic suppression. This effect is commonly studied with large, often peripheral, stimuli presented during instructed saccades. However, little is known about how saccades modulate the foveola and how the resulting dynamics unfold during natural visual exploration. Here we measured the foveal dynamics of saccadic suppression in a naturalistic high-acuity task, a task designed after primate’s social grooming, which—like most explorations of fine patterns—primarily elicits minute saccades (microsaccades). Leveraging on recent advances in gaze-contingent display control, we were able to systematically map the peri-saccadic time-course of sensitivity across the foveola. We show that contrast sensitivity is not uniform across this region and that both the extent and dynamics of saccadic suppression vary within the foveola. Suppression is stronger and faster in the most central portion, where sensitivity is generally higher and selectively rebounds at the onset of a new fixation. These results shed new light on the modulations experienced by foveal vision during the saccade-fixation cycle and explain some of the benefits of microsaccades.

## Introduction

Human vision is not uniform across space. While the retina collects information from a broad field, only a minuscule fraction—less than 0.01%—is examined at high resolution. This is the area covered by the foveola, the region void of rods and capillaries, where cones are most densely packed. Because of this organization, rapid eye movements, known as saccades, are necessary to redirect gaze toward the objects of interest, abruptly translating the image across the retina every few hundreds of milliseconds. It is remarkable that the visual system appears unperturbed by these sudden visual transitions and seamlessly integrates fixations into a stable representation of the visual scene.

It has long been observed that visual sensitivity is transiently attenuated around the time of saccades, a phenomenon believed to play a role in perceptually suppressing retinal image motion during eye movements. This effect, known as “saccadic suppression”, consists in the elevation of contrast thresholds to briefly flashed stimuli, which precedes the initiation of the saccade and outlasts it by as much as 100 ms^1–9^. Saccadic suppression is typically investigated with stimuli that cover large portions of the visual field, often in the periphery. Limitations in the precision of stimulus delivery, both spatial and temporal, have so far prevented mapping of the saccade-induced dynamics of visibility within the foveola. Thus, despite the disproportionate importance of foveal vision, little is currently known about its time-course around the time of saccades.

Studies on saccadic suppression also commonly focus on large saccades under well-controlled, but artificial, laboratory conditions. However, an examination of the time-course of foveal vision needs to take into account that natural execution of high-acuity tasks—the tasks that require foveal vision—normally tends to elicit saccades with very small amplitudes ^10^. Microsaccades, gaze shifts so small that the attended stimulus remains within the foveola, are the most frequent saccades when examining a distant face^11^, threading a needle^12^, or reading fine print^13^, tasks in which they shift the line of sight with surprising precision.

Because of their minute amplitudes, microsaccades pose specific challenges to the mechanisms traditionally held responsible for saccadic suppression^14–22^. These movements yield broadly overlapping pre- and post-saccadic images within the fovea, which would appear to provide little masking in visual stimulation^23–25^. They also result in reduced retinal smear^26,27^, as they rotate the eye at much lower speeds than larger saccades, delivering luminance modulations that are well within the range of human temporal sensitivity. Furthermore, it is unknown whether possible corollary discharges associated with microsaccades exert similar effects to their larger counterparts^28,29^. Despite these observations, microsaccades have been found to generally suppress vision^30–33^. However, the only two studies that specifically examined foveal vision during microsaccades reached diametrically opposite conclusions, with one arguing for a normal reduction in sensitivity^34^ and the other for a complete lack of suppression^35^.

Mapping peri-saccadic visual dynamics across the foveola is technically challenging. An immediate difficulty comes from targeting stimulation to a specific location within a region this small, since that the entire foveola is comparable in size to the region of uncertainty in gaze localization resulting from standard eye-tracking methods. This is further complicated by the very high speeds and brief durations of saccades, which add the need for very precise timing of stimulus delivery to the already taxing requirements of spatial precision. However, recent advances in methods for gaze-contingent display control now enable determination of the line of sight with accuracy sufficient to selectively test selected foveal regions during normal eye movements. Leveraging on these recent advances, here we examined how microsaccades modulate foveal sensitivity during natural visual exploration.

We developed a gaze-contingent high-acuity task that resembles primate social grooming, a task that very naturally integrates visual search and detection of brief stimuli and that spontaneously elicits frequent microsaccades. By precisely tracking gaze and presenting probes at desired retinal locations with high spatial and temporal resolution, we were able to map the peri-saccadic dynamics of contrast sensitivity across the foveola. Our results show that microsaccades are accompanied by an elevation of visual thresholds at the center of gaze that starts before the initiation of the movement but dissipates very rapidly as the saccade ends. The extent and dynamics of this suppression vary with eccentricity across the foveola, so that a stronger modulation occurs in the most central region, where vision is selectively enhanced after a saccade.

## Methods

### Subjects

Data were collected from six subjects (5 females and one male; age range 25-33 years). To ensure high quality of eye tracking and gaze-contingent display control, only emmetropic observers with at least 20/20 acuity, as tested with a standard eye-chart exam, were allowed to participate. With the exception of one of the authors, all observers were naïve about the purposes of the experiments and were compensated for their participation. Informed consent was obtained from all participants following the procedures approved by Boston University Charles River Campus Institutional Review Board and the Declaration of Helsinki.

### Apparatus

Stimuli were displayed on a fast-phosphor calibrated CRT display (Iyamaya HM204DT) at a resolution of 800 × 600 pixels and a refresh rate of 200 Hz. Observers were maintained at a fixed distance from the display, so that each pixel on the monitor subtended an angle of ~1.3’. Movements of the head were minimized by means of a head-rest and a custom dental-imprint bite-bar. Stimuli were viewed monocularly with the right eye while the left eye was patched.

Eye movements were recorded by means of a Generation 6 Dual Purkinje Image (DPI) eye-tracker (Fourward Technologies). The internal noise of this device has root mean square smaller than 20 seconds of arc^36^, enabling measurement of eye movements with approximately 1 minute of arc resolution as assessed by means of an artificial eye^37^. Vertical and horizontal eye positions were first low-pass filtered (cut-off frequency 500 Hz) and then sampled at 1 kHz.

### Stimuli

Stimuli were designed to loosely replicate the visual input signals experienced by primates while engaged in grooming. They consisted of 30 dark dots (the test locations; each a 5’ dark square at 2.8 cd/m^2^ luminance) simulating “fleas” and “dust particles”, which were randomly distributed within the central 2° region of the display. These objects were displayed over a naturalistic noise field background, which simulated the “fur” of the animal and covered the entire display, approximately 17° of visual angle. The power spectrum of the background decreased proportionally to the square of the spatial frequency as it happens in natural scenes^38^. The average luminance of the display was ~7 cd/m^2^. An example of the central portion of the stimulus in a trial—the region containing the dots—is shown in Fig. 1*A*. A different noise pattern and array of dots were displayed on each trial. Stimuli were rendered in OpenGL and modified in real-time according to the observer’s eye movements using EyeRIS, a custom system for gaze-contingent display control^39^. This system is designed to guarantee precise timing between changes in the stimulus and oculomotor events (typical delay 7.5 ms) and has been tested extensively see supplementary information in^10,40^.

**Figure 1:**
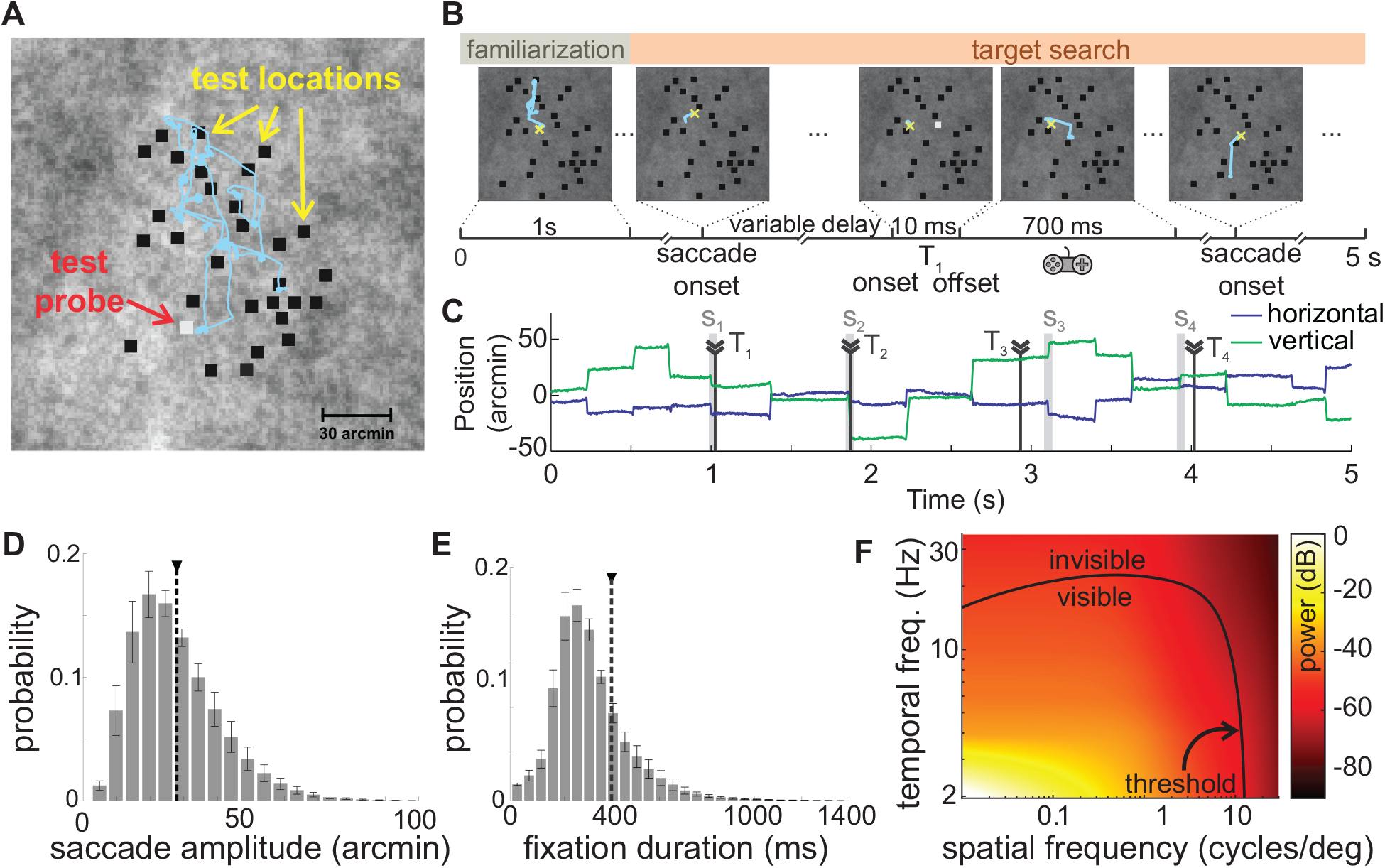
A virtual grooming task. Observers were instructed to search for fleas hiding within the animal’s fur (a naturalistic noise field). (**A**) 30 dots (5′ width) were distributed at random test locations across the central 2° region of the display. Subjects were told that a few of these dots were fleas and would distinguish themselves from the remaining “dust particles” by occasionally jumping (a 10 ms contrast pulse; the probe). The subject’s goal was to “catch” each flea as soon as it jumped by pressing a button on a joypad. (**B-D**) Example of a trial. (**B**) Following an initial familiarization period (1 s), the onset of a saccade triggered, with variable delay, a probe (*T_k_*) at one of the test locations. Both location and timing were selected in real-time according to the observer’s eye movements to test performance at various positions in the fovea and lags relative to saccades. The yellow cross and cyan segments represents the center of gaze and eye movements, respectively. (**C**) Gaze position during the course of the trial. Each probe was associated with the closest saccade (*S_k_* in *C*). Only probes with no more than one saccade within a ±200 ms window were selected for data analysis. (**D-E**) Characteristics of eye movements. Average distributions of saccade amplitude (*D*) and intersaccadic intervals (*E*) across *N* =6 observers. Error bars represent SEM. Vertical dashed lines mark the means of the distributions. (**F**) Power spectrum of the luminance flow delivered to the retina by the recorded saccades. The black line marks contrast sensitivity thresholds in humans (data from Kelly ^46^). The small saccades recorded in this experiment yield visual signals well within the ranges of spatiotemporal sensitivity.

### Procedure

Data were collected in separate sessions, each lasting approximately one hour. Every session started with preliminary procedures to ensure optimal eye tracking and gaze-contingent control. Data were then collected in blocks lasting 10-15 minutes, with breaks between blocks to allow the subject to rest. Every experimental session consisted of 5 blocks of 40 trials.

Accurate localization of the line of sight was achieved by means of a gaze-contingent two-step calibration already described in previous publications^10,41^. During the first stage of this procedure, subjects completed a standard 9-point calibration by sequentially looking at markers of a 3×3 grid. In the second stage, observers used a joypad to finely refine the estimated location of the center of gaze, which was displayed in real-time on the monitor. This refinement was also repeated after every trial for the central point of the grid to compensate for possible drifts in the apparatus and/or small head adjustments that may also occur under head immobilization. We have previously shown that this gaze-contingent calibration improves localization of the line of sight by approximately one order of magnitude over standard methods^10^.

To measure contrast sensitivity during normal oculomotor activity, we developed a “grooming task”, a high-acuity task designed after primate’s social grooming that naturally incorporates visual search and detection of transient events (Fig. 1*B*). Observers were instructed to search for fleas (dark dots) hidden within the fur of an animal (the noise field). They were told that some of the dots at the test locations were fleas whereas others were dust particles, and that the fleas would occasionally reveal themselves by “jumping”, a 10 ms pulse in luminance (the probe) that randomly occurred during the trial. Observers were asked to catch each flea as soon as they saw it jumping by pressing a button on a joypad.

To assess sensitivity at various retinal positions and different times relative to saccades, the observer’s eye movements were continually monitored, and the probes delivered in a gaze-contingent fashion timed to the onset of saccades, as signaled in real-time by EyeRIS (estimated instantaneous speed > 9°/s). Following detection of a saccade, one of the test locations was selected and the probe activated after a random delay (0-400 ms). Selection of the probe location was based on the current location of gaze, so to uniformly sample the two quadrants on the retina centered on the horizontal meridian at eccentricities smaller than 1° (Fig. 2A). To test one specific retinal location and temporal delay, a saccade could only activate one probe, and each probe was followed by a 700 ms refractory period during which no other contrast pulse occurred. Precise timing of all relevant events was saved by EyeRIS for offline analysis. Subjects were not informed of any of the rules determining the presentation of the probes and remained fully unaware that changes in the display were triggered by their eye movements.

**Figure 2:**
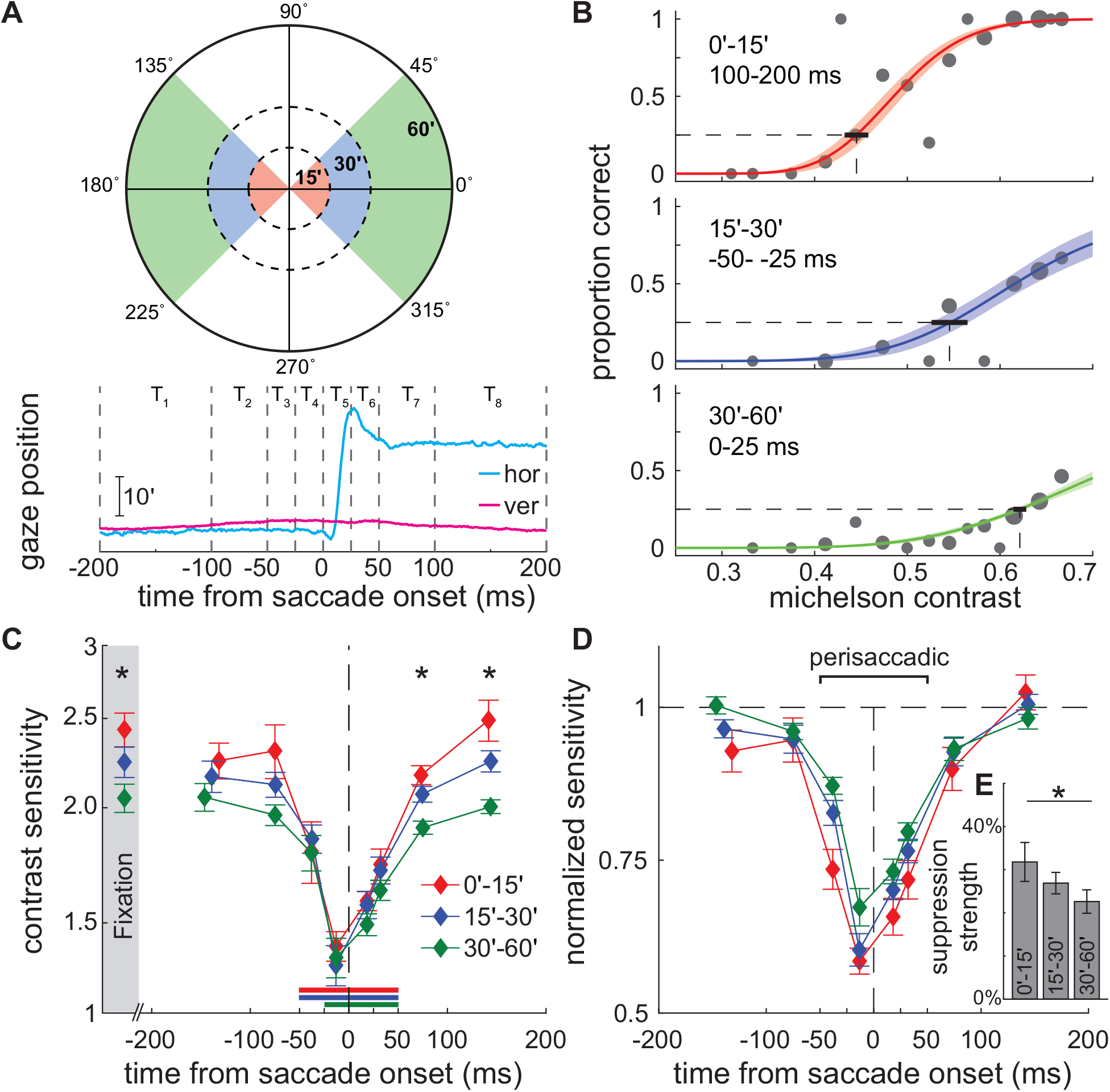
Changes in foveal sensitivity at the time of saccades. (**A**) Contrast sensitivity was measured in 24 spatiotemporal bins around saccades: 3 distinct regions within the fovea (eccentricity 0-15′, 15-30′, and 30-60′; top panel); and 8 time intervals around a saccade (bottom). (**B**) Contrast sensitivity functions in three spatiotemporal intervals for one observer. Colored lines and shaded regions represent, respectively, the maximum likelihood fitting and its SEM of a cumulative log-normal function to the data (gray circles; size proportional to the number of samples). The thick horizontal segment represents the SEM of the estimated 25% threshold. (**C**) Dynamics of contrast sensitivity relative to saccade onset. Each line represent mean sensitivity across observers (*N* = 6) in a foveal region. Error bars are SEMs. For comparison, sensitivity measured at fixation, when the probe appeared at saccade lags larger than 200 ms is also shown (shaded region). Horizontal bars indicate the intervals in which sensitivity deviated significantly from fixation (*p* < 0.05, post-hoc Tukey-Kramer comparisons). ⋆ marks significant differences across foveal regions (p < 0.05, one-way ANOVA). **(D**) The same data after normalizing each foveal region by its sensitivity at fixation to highlight differences in dynamics. (**E**) Mean perisaccadic suppression strength across the foveola. Suppression is strongest in the central region (p < 0.05, post-hoc Tukey-Kramer comparisons).

Every trial had a fixed duration of 5 s. It started with the presentation of a new stimulus (a new noise field and pattern of dots) and consisted of two phases: familiarization and search (Fig 1B). No probe was activated during the initial 1-s familiarization phase. Probes were displayed during the search phase and marked as hits if they were followed by a button press within 0.3-1 s, an interval chosen based on the typical range of reaction times. In each trial, the change in luminance (the amplitude of the pulse) varied randomly among 20 possible values ranging from 2.8 to 11 cd/m^2^, with the latter corresponding to the maximum intensity allowed by the monitor’s settings. Luminance steps varied occasionally across experimental sessions to ensure accurate fitting of the psychometric functions.

The number of probes in a trial varied depending on the numbers of saccades performed by the observer. Subjects were run extensively to estimate contrast sensitivity functions at 3 retinal eccentricities and 8 times relative to saccades. On average, 17,000 probes in 6,000 trials were collected from each observer in approximately 30 experimental sessions.

### Data analysis

Recorded oculomotor traces and the events data saved by EyeRIS were examined offline to determine the precise position of each probe on the retina and its timing relative to saccades. Only blink-free trials with optimal, uninterrupted eye-tracking were selected for analysis.

We first segmented each trace into complementary periods of saccades and fixations based on the eye speed. Data segments in which the eye displaced by more than 3’ reaching a speed of 3°/s were marked as possible saccades, and their onset and offset defined as the initial and final times at which the eye speed exceeded and returned below 2°/s, respectively. Consecutive events closer than 15 ms were merged together, a method that automatically takes care of possible post-saccadic overshoots^42^. Segmentation of the traces was performed automatically and then validated by manual inspection. Saccade amplitude was defined as the modulus of the vector connecting the eye positions at onset and offset.

Each probe was associated with the closest saccade based on temporal proximity. This event was not necessarily the one that triggered the probe during the course of the trial, as other saccades could have occurred closer to the probe, as in the examples S3 and S4 in Fig. 1*C*. Only probes with no more than one saccade within ±200 ms and associated with saccades smaller than 1° were considered in the analyses. To examine visual sensitivity at various visual eccentricities and lags relative to saccades, probes were clustered in 24 spatiotemporal bins (Fig. 2*A*). In space, we mapped performance in three eccentricity ranges in the ±45° circular sectors centered on the horizontal median: 0-15’, 15-30’ and 30-60’. In time, we examined the evolution of contrast sensitivity at 8 intervals around selected saccadic events: saccade onset, offset and peak velocity.

Psychometric functions of contrast sensitivity were independently evaluated in each of these bins via a maximum likelihood procedure (examples in Fig. 2*B*). A cumulative log-normal function was fitted to the performance data measured at various contrasts by means of the Psignifit Matlab toolbox^43,44^. The contrast sensitivity values reported in Figs. 2-5 are the inverse of the Michelson contrast:

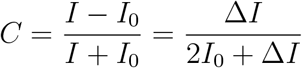

where *I*_0_ represents the baseline intensity of the probe (~3 cd/m^2^) and Δ*I* is the change in luminance of the pulse (*I* = *I*_0_ + Δ*I*). Virtually identical results were obtained by measuring changes in Weber contrast of the probe relative to its surroundings (Supplementary Fig. S2). To follow the dynamics of the low visibility measured around the time of saccades, we summarized performance by the contrast thresholds yielding 25% correct detection. For each subject and spatiotemporal bin, variability in the estimated threshold was assessed by non-parametric bootstrap over 1,000 random samples of the probes (error bars in Supplementary Fig. S1). Subjects were run extensively to collect sufficient amount of data for a reliable estimation of contrast thresholds in all bins. On average, 10,000 probes were used to construct the spatiotemporal map of contrast sensitivity for each individual, corresponding to ~65% of the total number of probes.

The spectral density map in Fig. 1*F* was obtained by reconstructing the luminance signals delivered by the recorded saccades and estimating their power spectra. The spectral analysis was conducted using an approach that allows high spatial resolution, as previously described in the literature^45^, and then averaged across subjects. The lines mapping the range of visibility are typical contrast thresholds taken from classical measurements of human sensitivity in the absence of retinal image motion^46^.

## Results

In a simulated grooming task, observers reported the occurrence of ‘flea jumps’ (the probes), brief changes in the luminance of otherwise dark dots located within the central 2° region of a wide naturalistic noise field. Subjects freely moved their eyes searching for the locations at which these contrast pulses would occur, while their eye movements were continually recorded.

In reality, unbeknownst to the subject, the probes were activated on the basis of the position and movement of the eye to measure visibility within selected regions of the fovea and at various time lags around saccades. This was possible thanks to three state-of-the-art components: (1) high resolution eye-tracking achieved via the Dual Purkinje image method^36^; (2) accurate gaze localization obtained by means of an iterative gaze-contingent calibration, a procedure that improves accuracy by approximately one order of magnitude over standard methods^10^; and (3) real-time control of retinal stimulation, obtained via a custom system for flexible gaze-contingent display control EyeRIS;^39^.

As expected, this high-acuity task resulted in the frequent occurrence of minute saccades. On average, observers executed ~2.5 saccades/s, almost all of them smaller than 1° (average saccade amplitude and standard deviation across subjects: 28’ ± 6’; mean 99th percentile of the amplitude distributions: 68’; Fig. 1*E*). In fact, the majority of saccades (68%) were smaller than 30’, an amplitude range that maintains an initially foveated probe well within the foveola—the microsaccade range, as defined in [10]. These small gaze shifts occurred at a rate (1.6 microsaccades/s) much higher than those normally encountered in tasks that do not involve high visual acuity (typically < 0.2 microsaccade/s, *e.g.*, [47,48]), an observation consistent with the notion that microsaccades are normally part of the strategy for examining fine spatial detail^12,41,49^.

Because of their small amplitudes and stereotypical dynamics^45,50^, the saccades performed in this task resulted in relatively slow changes in visual stimulation. This, combined with the characteristics of the visual scene, which, like natural images, possessed predominant power at low spatial frequencies, resulted in luminance signals to the retina that were well within the range of human temporal sensitivity (Fig. 1*F*). Yet, as it happens for larger saccades, subjects were not aware of the resulting translations of the images on their retinas—the well-known phenomenon of saccadic omission^7^.

To quantitatively examine the consequences of saccades on foveal sensitivity, we binned contrast pulses according to their combinations of retinal locations and lags relative to saccade occurrence and separately estimated contrast sensitivity in each spatiotemporal interval (Fig. 2*A*). Fig. 2*B* shows the psychometric functions of contrast sensitivity measured for one subject in 3 spatiotemporal bins. As these examples show, sensitivity varied considerably not only with the timing of the probe relative to saccades, but also with its position on the retina, reaching, in some instances, low values even at the highest possible contrast.

We first examined sensitivity far from saccades. The data points in the shaded region in Fig. 2*C* represent the average thresholds across observers estimated during fixation, *i.e.*, when no saccade occurred in the surrounding ±200 ms of a probe. Strikingly, despite being separated by just a few arcminutes, the three considered foveal regions exhibited marked differences in sensitivity. Contrast sensitivity was always larger at the very center of gaze, and decreased with increasing eccentricity, so that sensitivity in the most central region (the region within 15’) was on average 8% higher than in the range 15-30’, which was in turn ~9% higher than sensitivity in the range 30-60’ (one-way ANOVA, *F* (2, 17) = 4.8; *p* = 0.02). These measurements represent, to the best of our knowledge, the first mapping of contrast sensitivity across distinct locations of the foveola. They show that, contrary to its anatomical homogeneity, sensitivity is not uniform within this region: optimal sensitivity is restricted to a very narrow region around the center of gaze during normal fixation.

As the probe approaches the onset of a saccade, drastic changes in visual sensitivity occur. Sensitivity drops sharply from the fixation baseline starting approximately 50 ms before the saccade and continues to be affected up to ~100 ms after the saccade onset, a time at which the saccade has typically already ended (Fig. 2*C*). At all the considered foveal locations, suppression was strongest in the 25 ms interval immediately preceding the saccade, when sensitivity dropped by approximately 38% on average. The dynamics of this effect was highly stereotypical across subjects, all of whom individually exhibited a similar and statistically significant attenuation in sensitivity (*p* < 0.05, nonparametric bootstrap; individual subject data in Supplementary Fig. S1). Thus, the minute saccades performed in our experiment were accompanied by a strong attenuation in sensitivity throughout the foveola, an effect qualitatively similar to the saccadic suppression observed elsewhere in the retina for larger saccades.

While suppression occurred over the entire foveola, the extent and time-course of the process differed across foveal regions. All regions ended up with similar visibility levels at the peak of the suppression. However, since sensitivity in distinct regions started from different fixation baselines, the amplitude and speed of the process also varied, so that the change in sensitivity was larger and faster in the most central region of the foveola than at other locations. On average in the 100 ms interval centered at saccade onset, sensitivity was attenuated by 33% in the central region with eccentricity smaller than 15’, whereas it was only reduced by 23% in the 30 60’ region (*p* < 0.001; post-hoc Tukey-Kramer comparison; Fig. 2*E*). Thus, given the similar overall duration of the effect across the foveola, both suppression and recovery were faster at the very center of gaze than at larger eccentricities (Fig. 2*D*). These results were robust relative to the specific methods for data analysis. Very similar results were obtained by measuring sensitivity to changes in the Weber contrast of the probe relative to its surroundings rather than the Michelson contrast of the probe alone (see Supplementary Fig. S2).

To better examine the temporal evolution of saccadic suppression, we recomputed the time-course of sensitivity relative to two distinct temporal events, the end of a saccade and the time at which a saccade reaches its peak speed. Visibility recovers extremely rapidly following a saccade. On average across foveal regions, sensitivity has returned to about 90% of its pre-saccadic value less than 25 ms after the saccade ends and is fully restored within an additional 25 ms (Fig. 3*A*). This happens because suppression largely precedes the actual movement of the eye. Suppression is already recovering by the time a saccade is in mid-flight and has reached its peak velocity (Fig. 3*B*). Normalizing each foveal region by its initial sensitivity further emphasizes the different dynamics occurring at distinct eccentricities. Changes in sensitivity proceed faster in the central region (< 15’), yielding a greater change around 100-50 ms before saccade offset than at larger eccentricities (*p* = 0.021; post-hoc Tukey-Kramer comparison; Fig. 3*C*). As a consequence, sensitivity levels off approximately 25 ms earlier in this central region relative to the more peripheral foveola (Fig. 3*D*).

**Figure 3:**
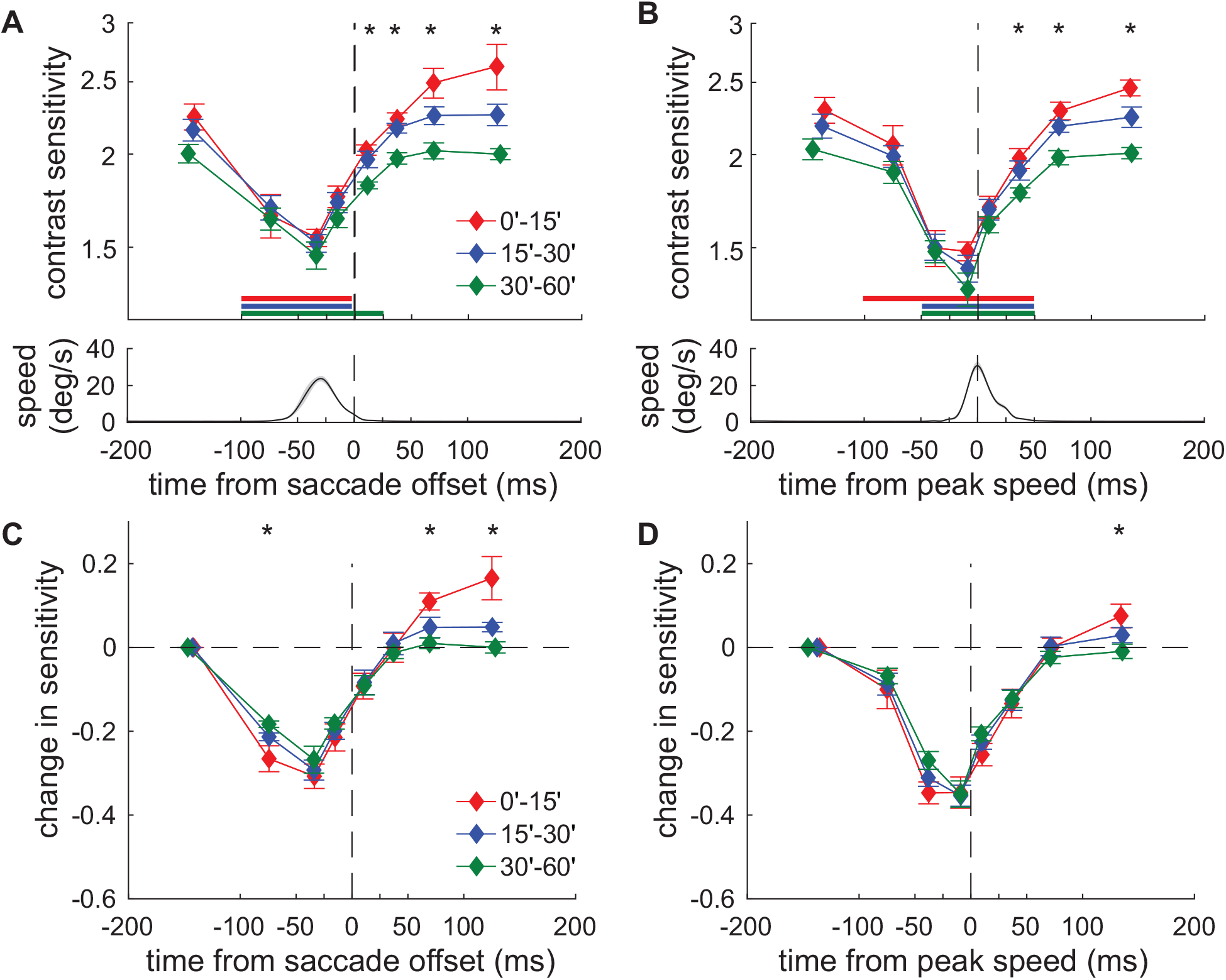
Dynamics of foveal sensitivity. (**A-B**) Average contrast thresholds are now aligned relative to either (*A*) the time at which the saccade ends, or (*B*) the time at which the saccade reaches its peak speed. Graphic conventions are as in Fig. 2*C* with the horizontal bars indicating statistically significant differences relative to fixation (*p* < 0.05, post-hoc Tukey-Kramer comparisons). The bottom panels show the mean instantaneous eye speed. (**C-D**) The same data normalized relative to the first sample to emphasize differences in dynamics across foveal regions. ⋆ marks significant differences across foveal regions (*p* < 0.05, one-way ANOVA).

The analyses in Figs. 2–3 were conducted including all saccades smaller than one degree. An interesting question is whether the time-course of saccadic suppression changes with the amplitude of the saccade, especially given that smaller saccades possess slower velocities and, therefore, maintain more stimulus power within the range of visibility on the retina. To investigate this question, we further subdivided the selected pool of saccades into two groups, depending on whether their amplitude was smaller or larger than 30’, and compared the dynamics of sensitivity elicited by each group. To counteract the loss in statistical power resulting from partitioning the saccade samples, we accumulated more data from each individual observer by comparing performance in two—rather than three—foveal regions, central and peripheral, depending on whether the probe was presented at eccentricities smaller or larger than 20’.

The outcome of this analysis is reported in Fig. 4, where the average temporal courses of sensitivity elicited by the two amplitude ranges are aligned relative to peak speed to enable clear comparison despite the different saccade sizes. The two groups of saccades had very similar effects on sensitivity. As already described in Figs. 2–3, modulations were sharper in the central portion of the foveola, where sensitivity started from higher values, and shallower in the peripheral region, where sensitivity was always lower. However, in both regions, the dynamics and extent of saccadic suppression were little influenced by saccade amplitude. Smaller saccades had virtually identical effects on visibility as larger ones, despite retaining more power within the range of human temporal sensitivity.

**Figure 4:**
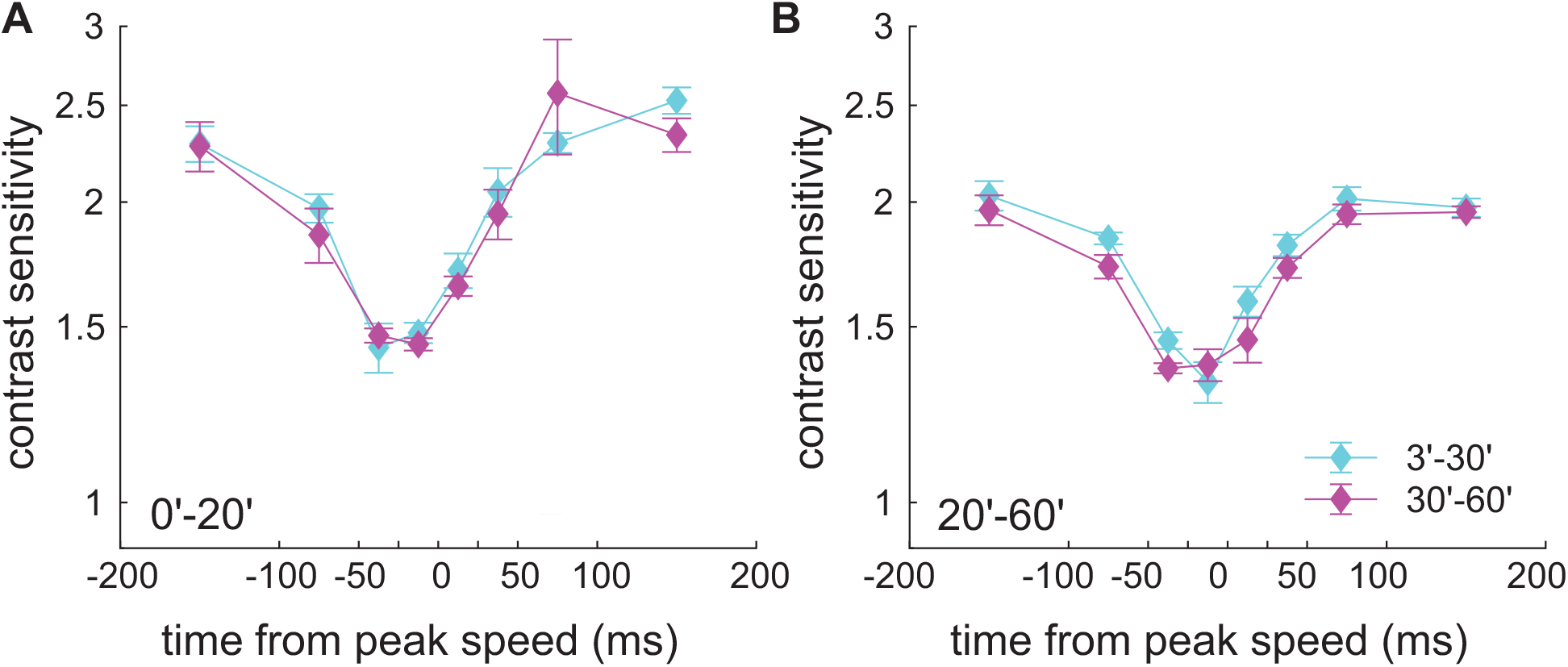
Foveal dynamics is unaffected by saccade amplitude. Changes in contrast sensitivity around saccades in distinct amplitude ranges: 0-30′ and 30-60′. Data represent averages across observers (N=6) aligned relative to the time of peak speed. Error bars are SEMs. The two panels show data in separate regions of the foveola: (**A**) the central region with eccentricity smaller than 20′; and (**B**) the more peripheral region at 20-60′.

Interestingly, sensitivity rebounds following a saccade, but only in the central portion of the foveola. This effect is clear in the data of Fig. 3*A*, which show that post-saccadic sensitivity continues to increase at the very center of gaze (< 15’) ~100 ms after a saccade, a time at which sensitivity has already saturated in more eccentric regions (*p* < 0.04; post-hoc Tukey-Kramer comparison). To examine in detail this post-saccadic enhancement, we directly compared levels of performance during pre-saccadic fixation, before suppression started, and in the fixation period that immediately followed a saccade.

Fig. 5 shows the result of this analysis for probes occurring 150 ms before saccade onset and in the 50-300 ms period after saccade offset. In the central foveola at eccentricity smaller than 20’, saccades were followed by significantly higher sensitivity, resulting in an average improvement across observers of 12% (p=0.027; paired two-tailed *t*-test). In contrast, sensitivity decreased following saccades in the more peripheral (≥ 20’) region of the foveola (p=0.308; paired two-tailed *t*-test). Thus, saccades led to opposite changes in sensitivity in these two regions (*p* = 0.001, paired two-tailed *t*-test). These results further emphasize the different modulations experienced by the distinct portions of the foveola in correspondence of saccades.

**Figure 5:**
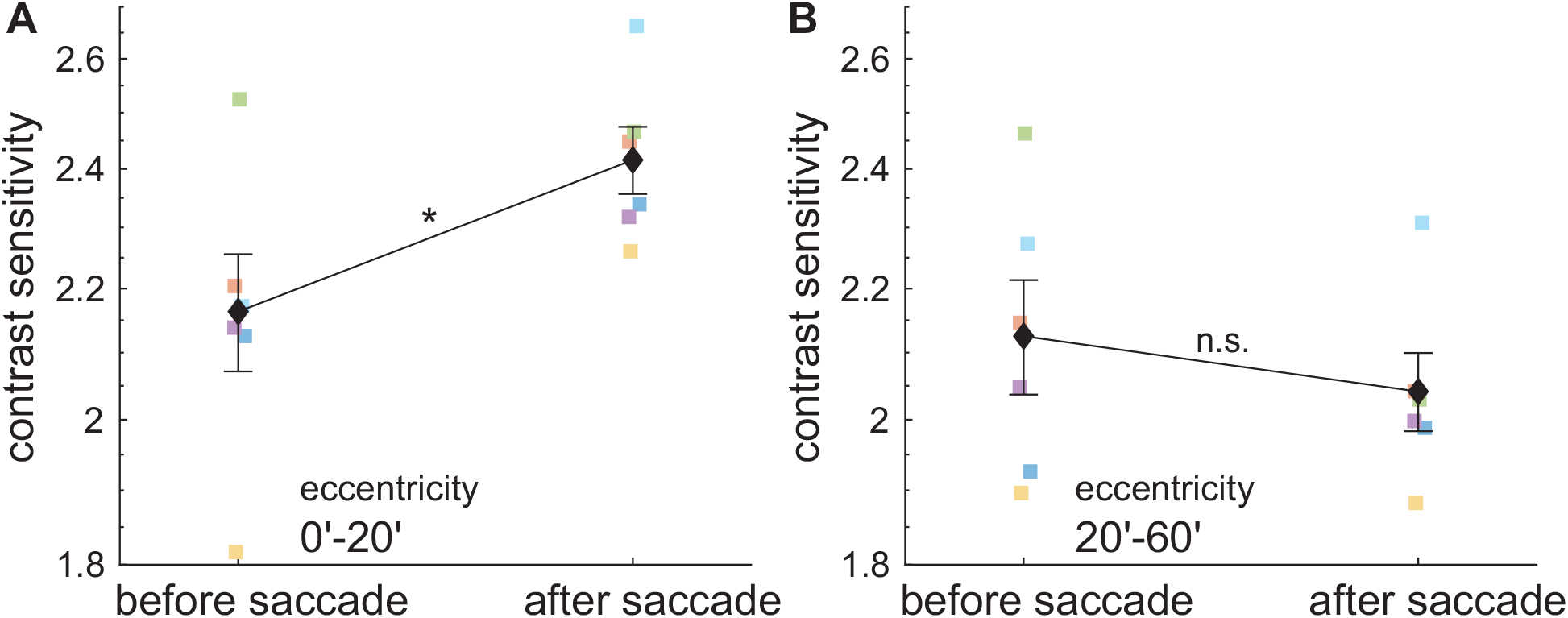
Post-saccadic enhancement is specific to the central foveola. Comparison of sensitivity before and after saccadic suppression. Black diamonds represent average sensitivity measured at least 150 ms before a saccade and 50-300 ms after. Error bars represent SEM. The two panels show data from: (**A**) the central foveola at eccentricity smaller than 20’; and (**B**) the peripheral foveola with eccentricity larger than 20’. Colored symbols are the individual subject data. ^⋆^*p* = 0.027, paired two-tailed *t*-test.

## Discussion

In this study, we examined the dynamics of foveal vision relative to the minute saccades that naturally emerge during fine spatial exploration. By implementing a naturalistic, yet highly controlled, high-acuity task, we were able to map contrast sensitivity at distinct locations within the foveola and follow their temporal evolution as eye movements occurred. Our results show that the foveola is accompanied by a general reduction in visual sensitivity in proximity of microsaccades. However, the extent and dynamics of this modulation are not uniform across the foveola: the attenuation is stronger and faster around the very center of gaze, where sensitivity rapidly rebounds at the end of the movement and remains higher than in the surrounding regions throughout post-saccadic fixation.

By providing the first mapping of contrast sensitivity and its peri-saccadic dynamics across the foveola, our results advance current knowledge of foveal vision in several ways. A first important finding is the observation that during normal inter-saccadic fixation, far from the occurrence of saccades, contrast sensitivity is not uniform in the central visual field, but varies by approximately 20%. This is a considerable change in such a small region and is consistent with previously reported impairments in discriminating small stimuli located slightly off the preferred retinal locus^41,51^. Previous studies did not measure contrast sensitivity across the fovea, but our results suggest that the way sensitivity declines with increasing eccentricity may have contributed to these impairments. It remains to be determined whether this attenuation in sensitivity originates from attentional modulations that transiently enhance performance at the very center of gaze or more sustained differences in neural processing with eccentricity. But irrespective of the specific mechanisms, our findings further highlight the importance for precisely controlling eye movements in tasks that involve fine spatial judgements, as the preferred retinal locus has a perceptual advantage relative to other foveal regions.

As the time of a saccade approaches, contrast sensitivity to briefly presented stimuli is attenuated throughout the foveola, a deficit that starts approximately 50 ms before the onset of the movement and dissipates very rapidly at saccade offset. As it happens at larger eccentricities, the time-course of the recovery is faster than that of the attenuation in all foveal regions, leading to significantly higher suppression in the initial part of the saccade, before reaching peak speed. Differences in modulations occur across foveal regions: since sensitivity is similarly impaired at the peak of the suppression, but its starting level depends on eccentricity, the amplitude and speed of the process varies across the foveola, so that the change in sensitivity is larger and faster in the most central region. These differences in dynamics are not captured by a simple multiplicative gain, as it has been proposed for larger saccades^52^, but represent more complex deviations in the shape of the modulations. They persist after normalizing each region by its pre-saccadic baseline to discount gain differences (Fig. 3). Even after normalization, sensitivity proceeds faster in the most central region. It deviates significantly from the other regions before the peak of the suppression (Fig. 3*C*) and levels off approximately 25 ms earlier (Fig. 3*D*). In addition, sensitivity rebounds in the central portion of the foveola, recovering faster and continuing to increase for longer than at larger eccentricities, an effect that leads to a considerable post-saccadic enhancement at the very center of gaze.

Qualitatively, these measurements resemble those previously reported outside the foveola, but important quantitative differences occur. Most evident is the overall strength of the effect, which is considerably weaker than the attenuation typically observed with larger saccades^1,15,52^—the commonly reported 0.5-1 log units suppression. On average across the foveola, sensitivity changed in our experiments by approximately half as much, 0.21 log units, a ~ 38% suppression. Multiple factors, including the small amplitudes of saccades and size of the probes, could have contributed to this reduction. However, it is important to point out that our naturalistic task differs substantially from the simplified stimuli often used by studies of saccade suppression. In these studies, probes are typically easily detectable when saccades do not occur. In contrast, in our experiments, the detection of probe was not always immediate even in the absence of saccades: the average values of sensitivity during fixation were such that it required 50% contrast modulation to reach threshold. Because of this, the moderate saccadic suppression measured in our experiments was sufficient to almost completely eliminate visibility of the probe around microsaccades.

Our work differs from previous investigations of visual sensitivity at the time of microsaccades in several important ways. A first fundamental difference is our focus on foveal vision. Most previous examinations used stimuli that covered large portion of the retina, often excluding the fovea^2,32,33,53^. Two notable exceptions that specifically focused on foveal vision reached opposite conclusions^34,35^, with the latter reporting lack of suppression during microsaccades. The reduced sensitivity measured in our experiments may help reconcile these previous findings, particularly in the light that—at least for larger movements^54^—suppression tends to be further attenuated for involuntary saccades. Importantly, no previous study has mapped the peri-saccadic time-course of sensitivity across the fovea, primarily because of the technical challenges inherent in the required spatiotemporal precision of retinal stimulation. These challenges were here overcome by leveraging on recent technological advances in gaze-contingent display^39^, which enabled coupling of high-resolution eye-tracking with accurate gaze localization and real-time control of stimulus delivery.

This study also differs from previous investigations for its focus on natural visual exploration. Studies on saccadic suppression typically focus on instructed saccades when dealing with larger movements and forced fixation when examining microsaccades. Both conditions occur rarely during natural viewing, when subjects typically react to stimuli by redirecting their gaze. Forced fixation, a condition in which observers are requested to maintain steady gaze on a marker, also creates a potentially problematic dissociation between the attentional demands of the motor task (the maintenance of fixation) and those of the perceptual task (the detection of a briefly presented stimulus), raising the possibility that disruptions and corrections for fixation mediated by microsaccades may temporarily distract from the visual task, transiently lowering detection performance. Here, we focused on the frequent tiny saccades that spontaneously emerge during normal examination of fine details. These movements are so small that they maintain the stimulus of interest well within the foveola, yet they carry perceptual benefits. Previous studies with accurate gaze localization have shown that, during natural exploration, microsaccades tend to precisely center the stimulus on task-relevant visual details^11–13^. Our present results show that this behavior would benefit from at least from two factors, the higher sensitivity around the preferred retinal locus and the post-saccadic sensitivity enhancement specific to this region.

An interesting perspective to the phenomenon of saccadic suppression comes from the proposal of active space-time encoding, the notion that luminance modulations from eye movements play a critical role in establishing spatial representations^55,56^. Intuition about the visual consequences of saccades is often gained by conceptualizing the resulting input signals as uniform (*i.e.*, constant-velocity) translations of the image on the retina (*e.g.* [57]). While much has been learned from this simplified model, saccades possess stereotypical dynamics^58^ that profoundly affect the luminance modulations delivered to the retina. Surprisingly, because of these dynamics, saccade modulations are qualitatively similar to those delivered by ocular drift, the incessant inter-saccadic motion of the eye^55^. On the retina, both saccades and drifts equalize (whiten) the power spectra of natural scenes within a low spatial frequency range. But a trade-off exists between power and bandwidth in this region: the larger a saccade, the narrower the whitening bandwidth and the higher the power it contains ^59^. These modulations are likely to play an important role in the strong responses elicited by the onset of a new fixation in the early visual system ^60–63^.

These considerations have important implications for saccadic suppression. First, the power delivered by saccades at low spatial frequencies is substantially lower than predicted from the standard uniform translation model. Thus, even a moderate suppression may be sufficient to prevent visibility of stationary scenes during saccades, a notion further reinforced by consideration that a brief flash is a particularly powerful stimulus. Second, one may expect stronger saccadic suppression with larger saccades, since the power of the visual flow in the whitening region increases with saccade amplitude. This prediction is consistent with previous findings^64^ as well with the relatively low level of suppression measured in our experiments. More broadly, the idea of active space-time encoding argues that, rather than being insensitive to the retinal image motion caused by saccades, the visual system uses these luminance modulations to encode visual information during post-saccadic fixation^59^ (see also [65–67]). The structure of these visual signals is consistent with the sensitivity enhancements at low spatial frequencies measured with saccades^68^, as well as with the dependence of this perceptual facilitation on saccade amplitude^45^.

In sum, our results show that contrast sensitivity is not uniform in the central fovea and is modulated by saccades in complex ways. Sensitivity is attenuated during microsaccades but recovers very rapidly and selectively rebounds at the very center of gaze. These modulations likely interact with task-dependent enhancements at specific retinal locations resulting from attentional shifts before the occurrence of saccades^69^ and microsaccades^70^. Further work is needed to investigate how these incessant foveal modulations influence oculomotor strategies and how humans actively deal with them to enhance visual performance.

## Data Availability

The data and matlab code to produce the main figures in the main text will be deposited into an online repository upon acceptance.

## Acknowledgements

This work was supported by the National Institutes of Health grants EY18363 (M.R.) and EY029565 (J.I.) and National Science Foundation grants 1457238 (M.R.). We thank Martina Poletti for helpful comments and discussion during the course of this research.

