## Supplemental Information for "Foveal vision at the time of microsaccades"

620 # Supplementary Information

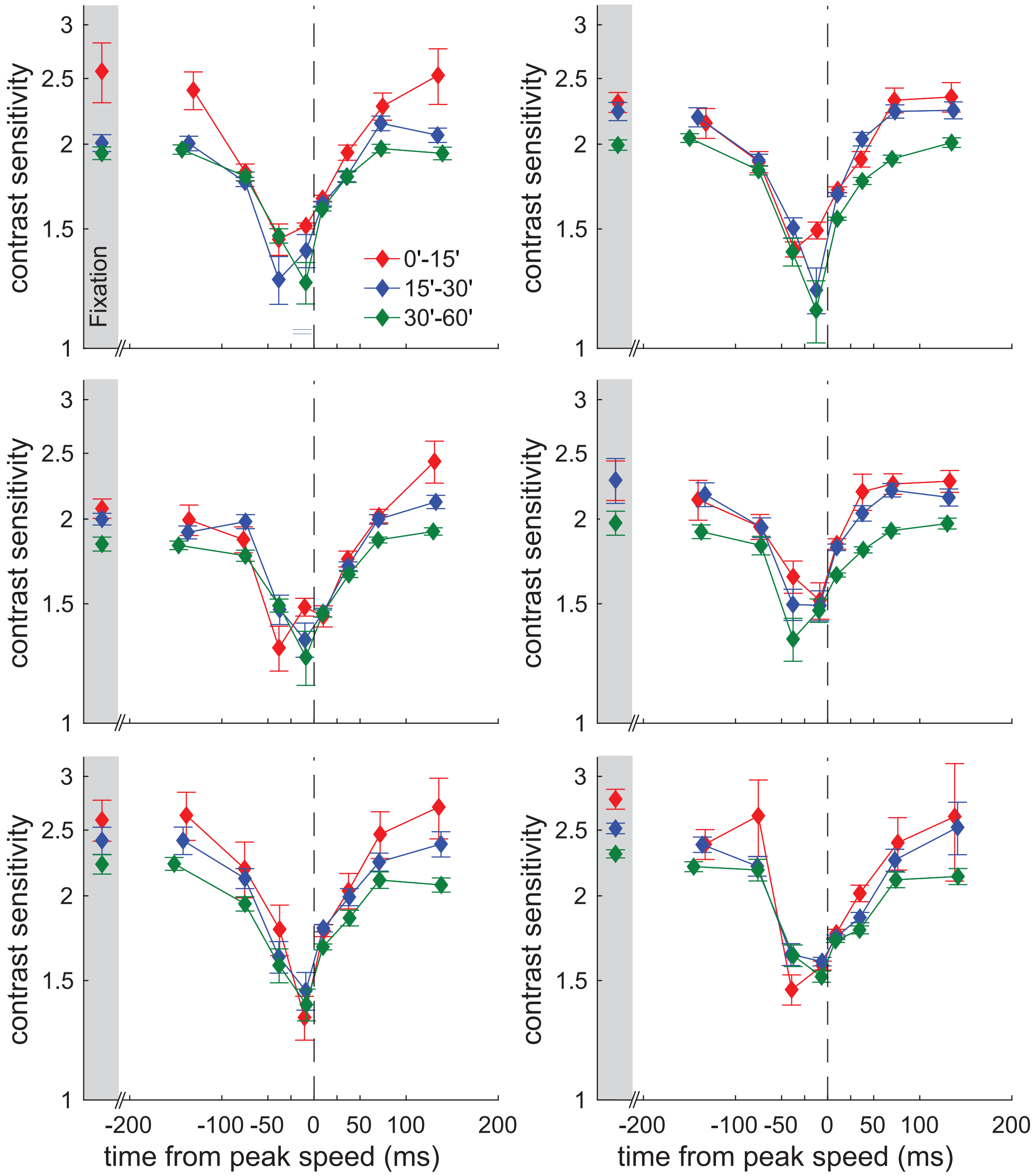


Figure S1: **Individual subjects' data.** Dynamics of contrast sensitivity relative to saccade peak speed. The data from the $N$=6 subjects are shows on separate panels. Graphics conventions are as in Fig. 2*C*. Error bars represent SEMs of the 25% contrast thresholds obtained from 1000 bootstrap repetitions.

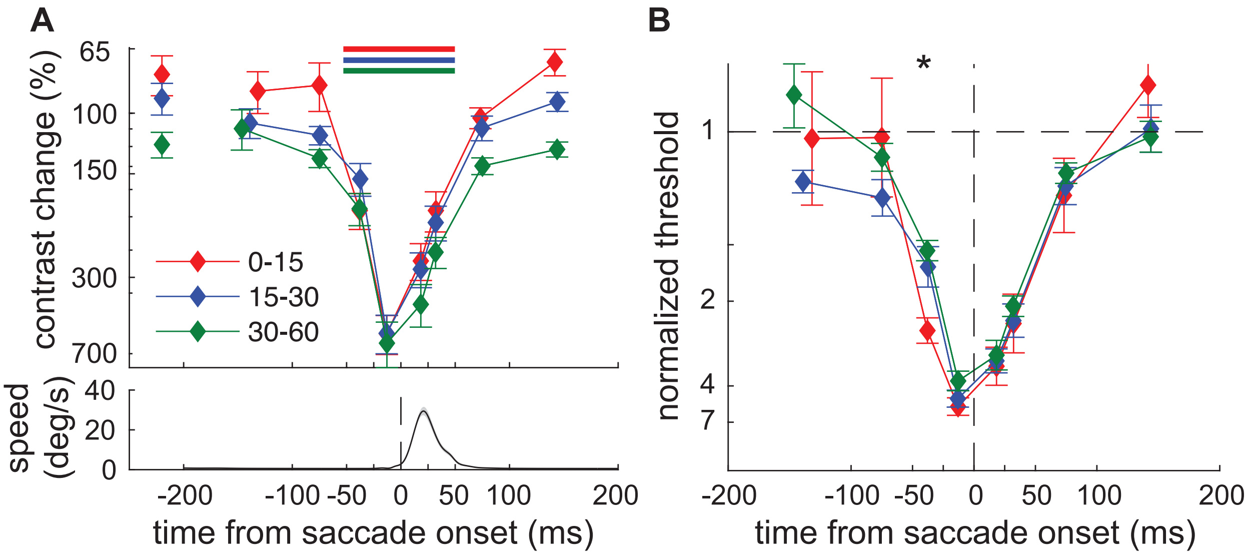


Figure S2: **Robustness of results.** Highly similar results were obtained by measuring sensitivity to changes in the Weber contrast of the probe relative to its neighborhood rather than the Michelson contrast of the probe alone. (**A**) Dynamics of sensitivity to Weber contrast changes relative to the time of saccade onset. Data represent 25% performance thresholds $(C_1 - C_0)/C_0$ averaged across subjects, where $C_0$ and $C_1$ are, respectively, the Weber contrasts of the dot before and after activating the probe relative to the average luminance of the display within the surrounding $10'$-radius circle. Horizontal bars indicate the intervals in which sensitivity differed significantly from the sample ~150 ms before saccade onset ($p < 0.05$, post-hoc Tukey-Kramer comparisons). (**B**) The same data normalized by the threshold measured at fixation. Values greater than 1 indicate visual suppression. Stars mark the intervals with statistically significant differences across eccentricities (one-way ANOVA, $F(2, 17) = 7.96$; $p = 0.004$).
